# Multiple Dynamic Modes of the Bcd Gradient are Explained by Quantum Mechanics

**DOI:** 10.1101/2024.04.04.588201

**Authors:** Irfan Lone, Carl O. Trindle

## Abstract

Extracellular diffusion coupled with degradation is considered as the dominant mechanism behind the establishment of morphogen gradients. However, the fundamental nature of these biophysical processes visa viz the Bicoid (Bcd) morphogen gradient remains unclear. Fluorescence correlation spectroscopy (FCS) has recently revealed multiple modes of Bcd transport at different spatial and temporal locations across the embryo. We here show that these observations, and a few others, are fitted by a model fundamentally based on quantum mechanics. We also indicate that the abstract and auxiliary feature called chirality of the said formalism finds a natural expression in our model of the Bcd gradient formation that might be verified in future experiments on the system.

## Introduction

Morphogen gradients play a crucial role in the early embryonic development of multicellular organisms by providing positional information to cells [1–5]. A prototypical morphogen gradient is that of the transcription factor Bicoid (Bcd) formed in the early Drosophila melanogaster (fruit fly) embryo and plays a key role in the determination of embryonal axis of the organism [6]. Although the Bcd gradient has been studied for a very long time [6–9], the biophysical mechanisms behind its establishment are still not completely understood. In 1970, Francis Crick suggested that morphogen gradients could be a consequence of passive diffusion [10], frequently assumed to be the mechanism associated with the biological transport processes. Crick introduced a model in which there is a constant production of morphogen at one end and its spatially uniform degradation throughout the system setting up a concentration gradient in three-dimensional space [11]. The state of such a system can be described as a classical random walk process, involving both diffusion and degradation, through the use of following reaction–diffusion equation,

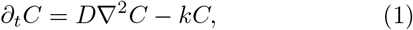

where *D* and *k* are, respectively, the diffusion and degradation constant of the morphogen, and *C* denotes its concentration. Further extensions of Crick’s ideas have essentially taken two directions. On the one hand the discovery of the Bcd gradient in the early fruit fly embryo [6] has motivated several diffusion-based models (see Ref. [12] and the references therein). On the other hand, fluorescence and single-molecule imaging based experiments [13–15] have inspired theories using explicit kinetic Markovian schemes based on the chemical master equation (reviewed in [16]):

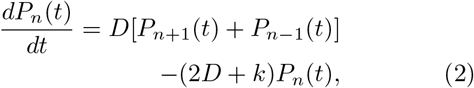

*for n >* 0, and

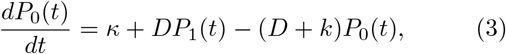

where *P*_*n*_(*t*) denotes the probability of finding the particle at the position *n* at time *t* and *κ* the morphogen production rate. In the steady-state regime, *dP*_*n*_(*t*)*/dt* = 0, and these differential equations simplify into a system of algebraic equations that can be solved analytically [16]. Further extensions accounting for both intrinsic noise [17–21] and diffusion [22–29] build upon Crick’s work. Regardless of the specific model details, all of these models share the common diffusion-degradation paradigm of Crick’s model in which the time evolution of the concentration fields, or probability densities, for the system is given by a deterministic reaction-diffusion equation or a coarse-grained approach based on above master equation.

In contrast, a recent experimental observation, utilising fluorescence correlation spectroscopy (FCS) and perturbative techniques, has revealed multiple modes of Bcd transport at different spatial and temporal locations across the embryo [30]. Exploring the different dynamic modes for Bcd it has been observed that in the cytoplasm the slow dynamic mode is similar across the embryo while as the fast one shows an increased diffusivity from the anterior to the posterior side [30]. Similarly, in the nuclei the Bcd diffusivity shows a significant, though comparatively small, increase in the fast dynamic mode from the anterior to the posterior regions, with the slow component remaining essentially unchanged [30].

The purpose of this Letter is to show that these observations, and a few others, can be explained through a quantum-classical treatment of Crick’s model. In our quantum-classical treatment, the degradation of Bcd is modelled as a unitary noise that is intrinsic and that does not cause any entanglement with the environment so that the system remains in an essentially pure state during the course of its evolution [31]. It thus acts like a classical fluctuating field such that the dynamics of the system is unitary, yet stochastic [32]. The mathematical mechanism proposed here is intrinsically different from that of the class of models developed previously. Thus, the state of the system at any given time in our model is represented by a vector Ψ in a finite-dimensional Hilbert space ℋ. Molecules are represented, just like in particle-based reaction–diffusion schemes [33], by point particles undergoing a Markovian quantum diffusion process in presence of a unitary noise that introduces a degree of stochasticity into their dynamics. Thus, our model is constructed in a bottom-up way based on the theory of quantum Markov processes and analytically solved for a one dimensional case [34]. We then show that the multiple dynamic modes of the Bcd gradient are a consequence of a quantum-classical dynamics. We conclude that the Bcd gradient is essentially a one-dimensional problem and thus a simple 1D quantum-classical model suffices for its formation dynamics.

## The quantum − classical model

Motivated in part by the experimental observation of multiple dynamics modes accompanying the Bcd gradient formation [30], we propose the model below (Fig. 1), which is equivalent to a modification of the Crick’s reaction-diffusion scheme, with degradation modelled as the unitary noise (Fig. 2). Compared to the usual classical random walk models, the coin toss is replaced by a chirality degree of freedom which can take two values denoted |+⟩ and |−⟩, for right- and left-handed chirality respectively [35]. A Bcd molecule of chirality state |+⟩ can move one step to the right at a time while that of chirality |−⟩ can move to the left along the lattice or they can also get degraded, leading to the setting up of a concentration gradient in one-dimensional space. Let ℋ_*n*_ be the position Hilbert space of the system. In our model ℋ_*n*_ has a support on the space of eigenfunctions |*n*⟩ corresponding to the sites *n* ∈ ℤ on the lattice, ℤ being a set of integers. This position Hilbert space ℋ_*n*_ is augmented by a ‘coin’ space ℋ_*c*_ spanned by the two basis states {|+⟩, |−⟩}. The states of the total system are thus in the Hilbert space which is the tensor product of above spaces: ℋ = ℋ _*n*_ ⊗ ℋ _*c*_ [36].

**FIG. 1.**
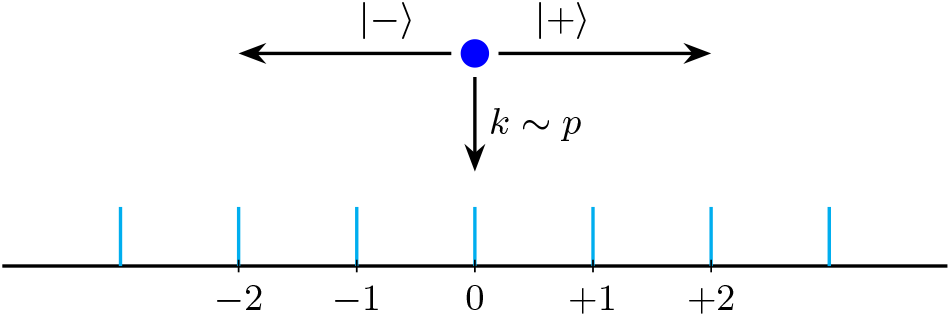
Schematic of the quantum-classical model. A Bcd molecule (shown by the blue dot) can, respectively, make a transition to the right or to the left depending on its chirality state of |+⟩ or |−⟩ or it can get degraded at a rate *k*.

**FIG. 2.**
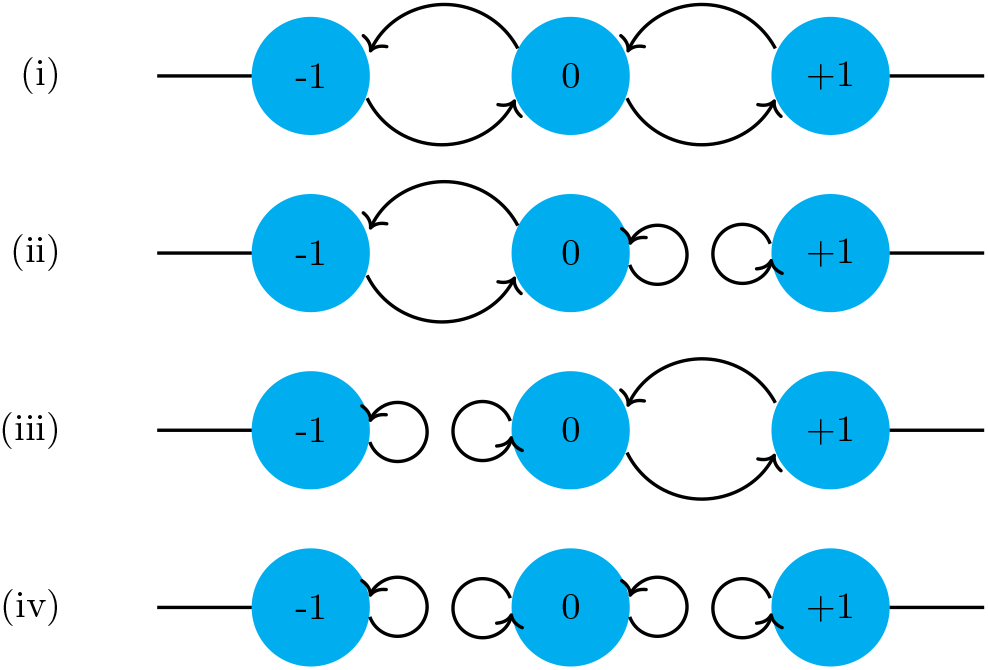
Schematic of the unitary noise in the Bcd system. The curved arrows depict the flow of probability flux between sites. Lattice sites, labelled by *n* = 0, ±1, ±2, …, correspond to the positions (states) of the Bcd particles in motion.

We now approximate the degradation of Bcd molecules by means of a unitary noise which proceeds as follows (Fig. 2). (i) When a Bcd molecule resides only briefly at a given site, it may not get internalized and degraded but instead survive and make a transition to the neighboring sites, |+⟩ to right and |−⟩ to the left. (ii) When a particle of chirality |+⟩ is internalized at a given site, e.g. *n* = 0, then there is no probability flux towards the nearby site, *n* = +1. Since the upper component of spinor at *n* = 0 sends probability flux towards *n* = − 1, in order to conserve flux, the outgoing probability flux from the lower component at *n* = 0 must be diverted to the upper component at the same site. Thus the transition that should have taken place is turned into a self-loop for that step of the walk. However, there is still a flow of probability flux towards the site *n* = − 1 due to the particles of |−⟩ chirality state. (iii) When a particle of chirality |−⟩ is internalized at a given site, *n* = 0 again, then there is no flow of probability flux towards the nearby site on the left, *n* = − 1, and the transition that should have taken place is again turned into a self-loop for that step of the walk. However, there is still a flow of probability flux towards the site *n* = +1 due to the particles of |+⟩ chirality state. (iv) When particles of both chirality states reside for appreciably longer periods of time at a given site, they may both get internalized and degraded. In such a situation there is no flow of probability flux in either direction. In this unitary noise model, we denote the probability per unit time of the Bcd molecule of a given chirality getting degraded by *p*. Thus, the parameter *p* quantifies noise in our model of the Bcd system.

We represent the quantum state of a Bcd molecule at a given time *t* by the following spinor

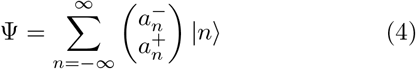

with the amplitude components 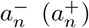 associated to the left (right) chirality states, respectively and | *n* ⟩ as the basis vector that locates the particle at *n* ∈ ℤ on the lattice with a discrete topology. The chirality consists of a single qubit (*a*^+^, *a*^−^)^*T*^. The probability of finding the walker at site *n* at a certain time *t* is given by

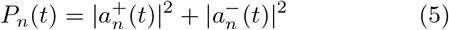

Now the particle moves along the integer lattice ℤ in discrete time steps depending on its coin state. The time evolution is expressed by

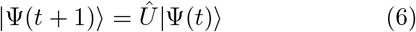

where 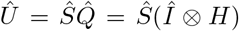 is a unitary operator that defines a step of the walk [37]. *H* is the Hadamard operator

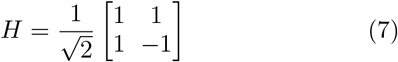

acting in the coin Hilbert space with the states of the chirality qubit expressed in the standard basis

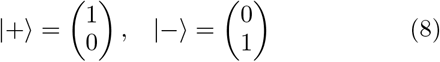

and *Î* the spatial identity operator [37]. These basis state vectors are transformed by the Hadamard operator as

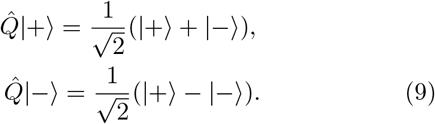

Next the conditional shift operator *Ŝ* translates the particle according to its chirality state as

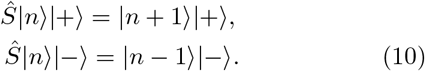

These operators together yield the desired recursion equations for the amplitudes of the quantum state [38].

There are four obvious cases to consider (see Fig. 2):

*Case a*. − this corresponds to the situation shown in Fig. 2(i) and is described by the following equation

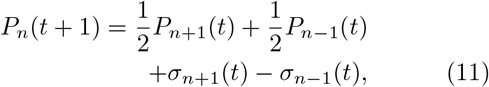

where *σ* ≡ ℜ [*a*^+*^*a*^−^] denote the quantum coherence terms and the symbol ℜ stands for the real part. Such, often fleeting, coherences can make some physical processes, like the ones important for quantum biology [39], more efficient by changing the scaling behavior of certain macroscopic observables [40].

*Case b*. − corresponds to the situation shown in Fig. 2(ii) and is represented by the following evolution equation

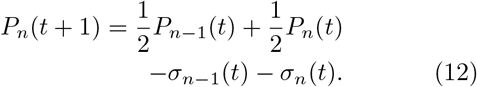

*Case c*. −this corresponds to the situation shown in Fig. 2(iii) and its evolution equation is given by

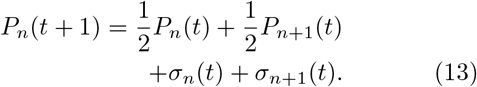

*Case d*. − finally this corresponds to the situation shown in Fig. 2(iv) and is represented by the following equation

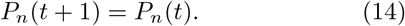

Eqs. (11-14) represent a set of quantum-classical evolution equations for our Bcd walker with the occupation probabilities *P*_*n*_(*t*) representing the classical part and the interference (*σ*) terms the non-classical contribution.

If we ignore the *σ* terms in the above set of quantum-classical equations and combine the resulting expressions into one single equation then a straightforward scaling argument gives the following result

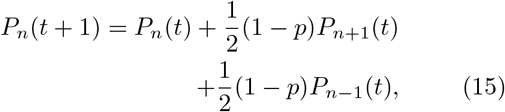

which yields the well-known diffusion equation

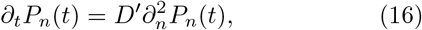

with the diffusion constant defined as

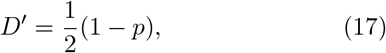

and (*n,t*) treated as continuous variables. Solution of the above set of quantum-classical Eqs. (11-14) without neglecting the coherence (*σ*) terms gives the following expression for the diffusion constant of the system

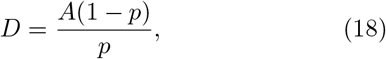

where the coefficient *A* is a function of *p* with an estimated value of ∼0.40 and *A* → 0.5 as *p* → 1 [41]. Remarkably, Crick, using his source-sink model,

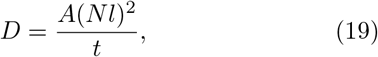

has estimated a value of *A* = 0.42 and proposed a general value of *A* = 0.5 for the biochemically more realistic models; where *t* is time, *N* is number of cells between source and sink, and *l* is length of each cell [10]. Equations (17) and (18) are used below to explain the origin of multiple dynamic modes that accompany the formation of the Bcd gradient in the early fruit fly syncytium.

### Multiple dynamic modes

Using florescence correlation spectroscopy (FCS) combined with certain perturbative techniques, Athilingam *et al*. have demonstrated that a two-component fit to the FCS curves is substantially better than a one-component fit in both the cytoplasm and nucleus [30], implying the presence of two dynamic modes in each region. There are thus, at least, four effective modes of Bcd transport: (1) fast cytoplasmic mode, (2) slow cytoplasmic mode, (3) fast nuclear localised mode, and (4) slow nuclear localised mode. The effective diffusion coefficient, *D*_*eff*_ = *g*_*f*_ *D*_*f*_ + *g*_*s*_*D*_*s*_, increases from the anterior to the posterior side of the embryo, where *D*_*f*_, *D*_*s*_ and *g*_*f*_, *g*_*s*_ represent the diffusion coefficients and fractions for the fast and slow dynamic modes respectively, with *g*_*f*_ + *f*_*s*_ = 1 [30]. Implementing such a spatial variation within a reaction-diffusion model, the authors of Ref. [30] have suggested that these multiple dynamic modes can generate a long-range yet steep gradient in a relatively short period of time. Overall, they have provided an improved version of the synthesis-diffusion-degradation (SDD) scheme for the formation of the Bcd gradient. But such a model leaves unanswered the question of the origin of multiple dynamic modes in the first place and their correlation with concentration.

Fig. (3) shows the plot of equation (17) as a function of noise, *p*. The variation of *D*^*′*^ versus *p* is linear and smooth with no large fluctuations as noise increases. Similar plot of Eq. (18) is shown in Fig. 4. As can be clearly seen from the graph, the diffusivity of the system is not constant as per this equation but varies with the magnitude of noise, reaching much higher values for very small noise levels. Clearly, in our model *D*_*s*_ and *D*_*f*_ correspond, respectively, to *D*^*′*^ and *D* from Eqs. (17) and (18). Therefore, the expression for the effective diffusivity of the system may also be expressed as

**FIG. 3.**
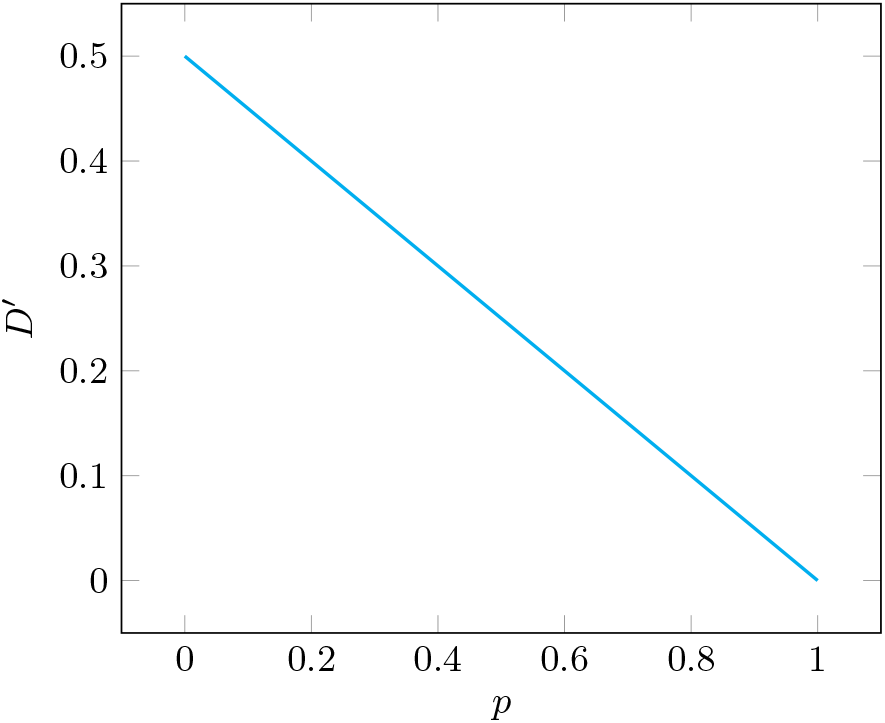
Slow dynamic mode of Bcd, equation (17), as a function of noise, *p*. There is no significant variation in *D*^*′*^.

**FIG. 4.**
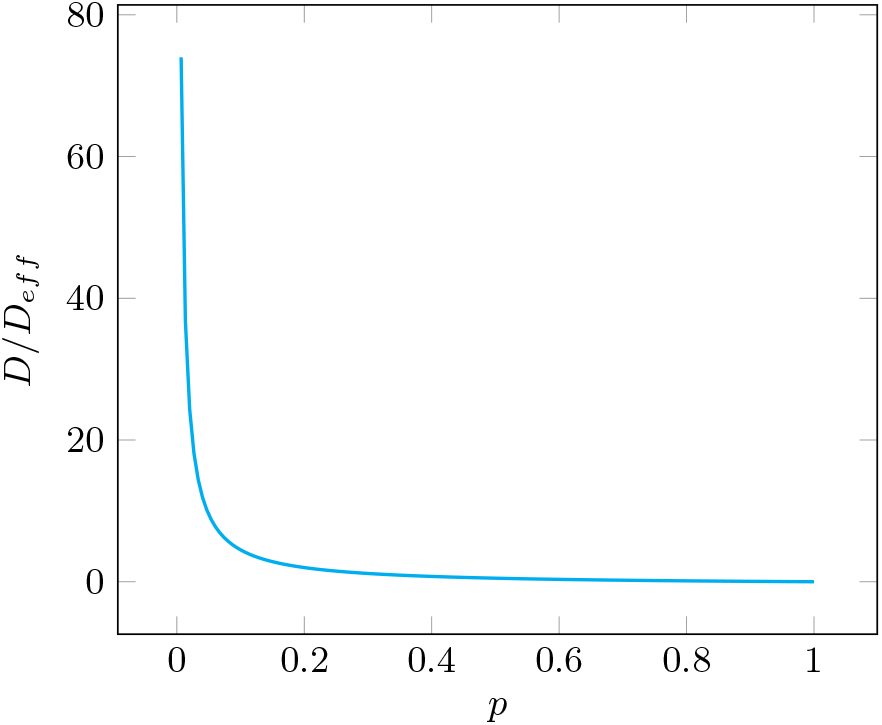
Fast dynamic mode of Bcd, equation (18), as a function of noise, *p*. The magnitude of *D* is large for very small *p*. Eq. (20) shows exactly similar behaviour as Eq. (18) for the cytoplasmic fractions of Bcd in both the anterior and posterior parts of the embryo.

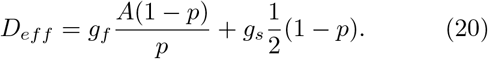

Equation (20) displays exactly similar behaviour as equation (18) for the cytoplasmic fractions of Bcd in both the anterior and posterior parts of the embryo with the values of *g*_*f*_ and *g*_*s*_ taken from Ref. [30]. This shows that the effective dynamics of the system is dominated by the dynamics of the faster mode in both the anterior and the posterior parts of the embryo. This is not surprising since the dynamics of the slower mode varies very little and remains nearly constant throughout the system in both the cytoplasm (*D*_*s*_ ∼ 1*μm*^2^*s*^−1^) as well as nuclei (*D*_*s*_ ∼ 0.3*μm*^2^*s*^−1^) [30]. This should also explain the rapid establishment of the Bcd gradient within a couple of hours [12]. Since a larger fraction of Bcd is confined to the nuclei during interphase [30], where most of the degradation and the actual transfer of positional information takes place [13], we shall confine our attention to the observed values of diffusivity for the nuclear dynamic modes. However, exactly the same analysis and equations will apply to the cytoplasmic modes. Now equation (17) is the same result as that for an unbiased classical random walk if we set *p* = 0. However, since *p* in reality is non-zero so the value of *D*^*′*^ should actually be less than 0.5 as per this equation. Experimental observations of Bcd dynamics have obtained the diffusivity values in the range of 0.2 − 0.3*μm*^2^*s*^−1^ for the slower dynamic modes in cycle 12-14 nuclei [30]. The Bcd gradient in the 500*μm* long syncytial embryo of Drosophila gets established in about 3 hours [12]. Within 2 hours the number of nuclei in the embryo increases by three orders of magnitude, from 1 to ∼ 6000. Taking the average nuclear diameter, *l* ∼10*μm* [12], the number of cells along the line between source and sink in the Crick’s model comes out to be about *N* ∼50. With the value of *A* = 0.5 and *t* = 3 hours, Eq. (19) gives *D* ∼ 12*μm*^2^*s*^−1^, which is of the same magnitude as that found for the effective diffusivity of Bcd experimentally [30]. Thus, even Crick’s source-sink model [10] is able to reproduce the observed dynamics of Bcd system, albeit without explaining the underlying microscopic mechanisms responsible for it.

Lattice light sheet microscopy studies have revealed highly dynamic Bcd molecules in the posterior end of the embryo with a diffusive-like kinetic behavior [15]. Recent FCS experiments have also measured a significant concentration of Bcd in the posterior-most parts of the embryo [30]. The required probability distribution for Bcd is calculated from *P*_*n*_(*t*) ≡ |*n* | Ψ(*t*) ⟩| ^2^. But one must first define an initial state for the system. The homochirality of amino acids ensures that Bcd, just like all other proteins, is a chiral biomolecule [42]. Since all naturally occurring amino acids that go on to constitute the proteins are left-handed, a reasonable initial state for Bcd is the one which is dominated by molecules of left-hand chirality conformation: |−⟩⊗|0⟩, where we assume the system to be in *n* = 0 state for the particles moving on an integer line. The probability distribution for Bcd is shown in Fig. (5) for N=1000 steps. Clearly it is steep and non-Guaussian with significant amplitude at both extremes and a vanishingly small amplitude towards the middle. The distribution thus predicts a small but significant concentration of morphogen on the posterior (righthand) side of the embryo. This explains the origin of a steep long-range gradient with a significant concentration of Bcd on the posterior side [30]. A remarkable fact is that such a distribution remains essentially unchanged for noise levels corresponding in magnitude (*p* ∼ 0.01) to observed Bcd degradation rates in the system [43].

At the core of our treatment lies the unitary coin operator, Eq. (7), that generates the requisite coherent superpositions of the chirality states. Quantum mechanics thus, in principle, allows for a coherent superposition of the left- and right-handed enantiomers of a chiral molecule. This chirality, which is otherwise an auxiliary degree of freedom introduced to implement a quantum Markov process, finds a natural expression in our model of the Bcd gradient dynamics. The phenomenon of Bcd gradient formation thus presents a potential opportunity for an experimental verification of these chirality superposition states since the existence of quantum superposition states for isolated biomolecular systems with masses (*>* 25*kDa*) in the range of Bcd (∼55*kDa*) is now already well established [44].

## Conclusion

This study is, to our knowledge, the first to employ the formalism of a quantum Markov process to analyze the dynamics of morphogen gradient formation. Using a hybrid model, we have shown that the multiple dynamic modes of the Bcd gradient are a consequence of quantum-classical dynamics. It remains to be seen whether a similar type of analysis can be carried out for other morphogen gradients, particularly the BMP gradient established orthogonal to Bcd along the dorso-ventral axis of the Drosophila melanogaster embryo [45].

The Bcd system is essentially a one-dimensional problem since patterning systems along different axes of the embryo are basically independent of each other [46]. Thus a simple one-dimensional quantum Markovian treatment becomes possible. Despite the apparent simplicity of our approach, it makes, in addition to explaining the observed features of the system, certain new and useful predictions.

We hope that our analysis of the Bcd gradient would provide a motivation for an exciting new line of experimental inquiry that may contribute to a richer understanding of the factors that play a decisive role during early embryonic development of multicellular organisms.

## Acknowledgements

We thank K N Tejas for executing the matlab code for Fig. 5. Dr Mir Faizal is acknowledged for encouragement.

**FIG. 5.**
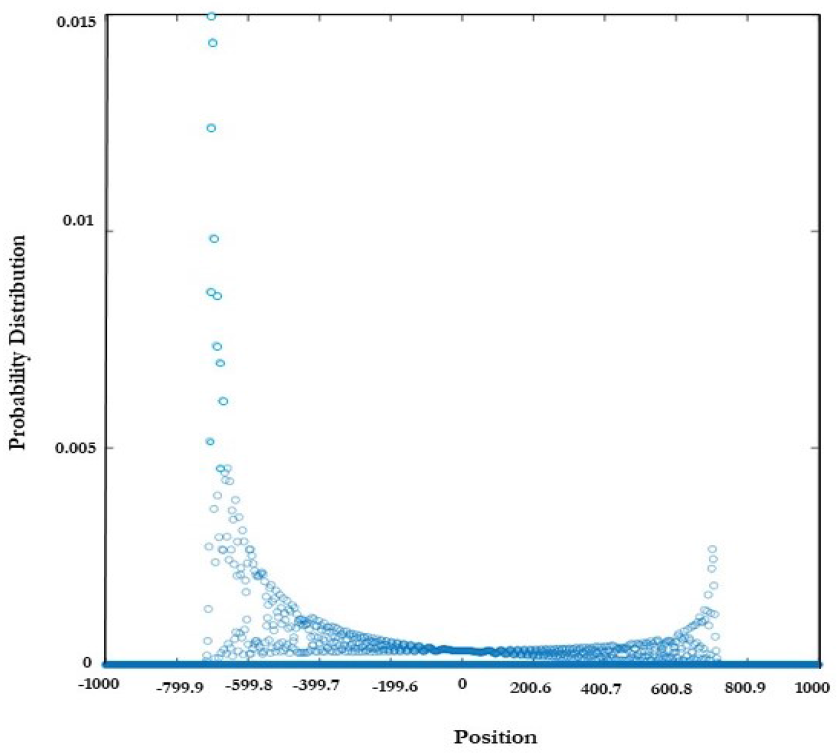
Probability distribution, *P*_*n*_(*t*) ≡ |⟨*n*|Ψ(*t*)⟩|^2^.

